# Range sizes of the world’s mammals, birds and amphibians from 5000 BCE to 2100 CE

**DOI:** 10.1101/779801

**Authors:** Robert M. Beyer, Andrea Manica

**Affiliations:** Department of Zoology, University of Cambridge, Downing Street, Cambridge CB2 3EJ, United Kingdom

## Abstract

Species’ vulnerability to extinction is strongly impacted by their geographical range size. Formulating effective conservation strategies therefore requires a better understanding of how the ranges of the world’s species have changed in the past, and how they will change under alternative future scenarios. Here, we used reconstructions of global land use and biomes of the past 7,000 years, and 16 possible climatic and socio-economic scenarios until the year 2100, to map the habitat ranges of 16,511 individual mammal, bird, and amphibian species through time. We estimate that extant species have lost an average of 19% of their natural range sizes thus far, and may lose up to 32% by 2100. Changes in range size vary greatly between species, with tropical, small-ranged, and threatened species being especially impacted. Our data reveal that range losses have been increasing disproportionately in relation to the size of destroyed habitat, driven by a long-term increase of land use in tropical biodiversity hotspots. The outcomes of different future land use and climate trajectories for global habitat ranges vary considerably, providing important quantitative evidence for conservation planners and policy makers of the costs and benefits of alternative pathways for the future of global biodiversity.

---

Habitat range size is a strong predictor of species’ vulnerability to extinction ^1,2^. As a result, two major drivers of the decline of geographic range sizes – the conversion of natural vegetation to agricultural and urban land, and the transformation of suitable habitat caused by to climate change – are considered the two most important threats to global terrestrial biodiversity ^3^. Land use change has caused staggering levels of habitat contractions for a range of mammal ^4,5^, bird ^6^ and amphibian species ^7^. Simultaneously, anthropogenic climate change has been driving shifts in species ranges ^8–11^, which, whilst resulting in larger range sizes for some species, has led to severe range retractions for others ^10,12,13^. Declines in global range sizes due to land use and climate change heavily contribute to the loss of local species richness ^14–16^ and abundance ^16–18^ in many parts of the world, thereby threatening essential ecosystem functions ^16,19^. With global agricultural area potentially experiencing a drastic increase in the coming decades ^20^, and climate change continuing to drive ecosystem change at an accelerating pace ^21^, future projections suggest that past trends in range contractions may continue ^22^, and likely contribute to projected large-scale faunal extinctions ^11,12,23,24^.

Considering the crucial role that species’ range sizes play for extinction risks, a better understanding of the long-term range dynamics of individual species, and projections of future changes under alternative scenarios, is crucial for conservation planning from the local to the global scale. Such estimates would allow quantification of historical pressures on species, and inform prioritisation of future efforts. Here, we estimated the habitat range sizes of 16,511 mammal, bird, and amphibian species since the early stages of agriculture until the end of this century based on global land use and climatic conditions. Accounting for these two factors simultaneously allows for the importance of interaction effects between them 25,26. We used empirical datasets on the global distribution of species, and combined these with species-specific biome preferences to estimate local habitat suitability under natural vegetation, cropland, pasture, and urban areas. By overlaying these data with reconstructions of global biomes corresponding to past climatic conditions, and agricultural and urban areas for the last 7,000 years, we estimated the historical habitat ranges of each individual species (Methods). We then extended the analysis into the future based on 16 alternative climate and land use trajectories until the year 2100, representing four emission scenarios (RCPs 2.6, 4.5, 6.0, 8.5), and five socio-economic pathways (SSPs 1–5) (Methods). SSPs represent different degrees of global socio-economic challenges for climate change mitigation and adaption: SSPs 1 and 4 correspond to small mitigation challenges, and SSPs 3 and 5 to high mitigation challenges; SSPs 1 and 5 correspond to high adaptation challenges, and SSPs 3 and 4 to high adaptation challenges; SSP 2 represents a middle-of-the-road scenario ^27,28^. RCPs 2.6–8.5 represent increasing levels of global warming by the end of the century ^29^. Considering all possible SSPs for any given RCP is crucial, as using only one realisation per RCP can conflate effects and lead to contestable patterns (e.g. ref. ^15^). We emphasise that our estimates of the geographic distribution of species’ habitat ranges based on land use and climate represent upper estimates for the actual distribution of populations. They do not account for other types of human influence, such as hunting ^30^, suppression by introduced species ^31^ and pathogens ^32^; nor do they incorporate the impacts of habitat fragmentation ^33^ and trophic cascade effects ^34^ on the viability of local populations.

With moderate impacts on the ranges of most species’ up until the industrial revolution, the expansion of agricultural production and settlements alongside the rise in population growth since the 18th century has drastically reduced range sizes of most mammals, birds, and amphibians (Fig. 1A). Using potential natural ranges in 1850 as a reference, we estimated that species had lost an average of 19% of their natural habitat by 2016. For most species, alterations in the global distribution of biomes due to past climatic change have had a much smaller effect on range sizes compared to land use, causing average range changes of less than 1% in the past 7,000 years (Fig. S1). There is substantial variability between species in terms of the experienced range changes. Species experiencing particularly severe range contractions include small-range species, and species classified as vulnerable and endangered (Fig. S2). Critical levels of range loss affect a rapidly rising number of species, with currently 15% having lost more than half of their natural range. Among these species, tropical species account for an increasingly larger proportion (Fig. 1B). For an estimated 18% of species, ranges have expanded in consequence of anthropogenic climate change and the conversion of unsuitable natural vegetation to cropland and pastures.

**Fig. 1.**
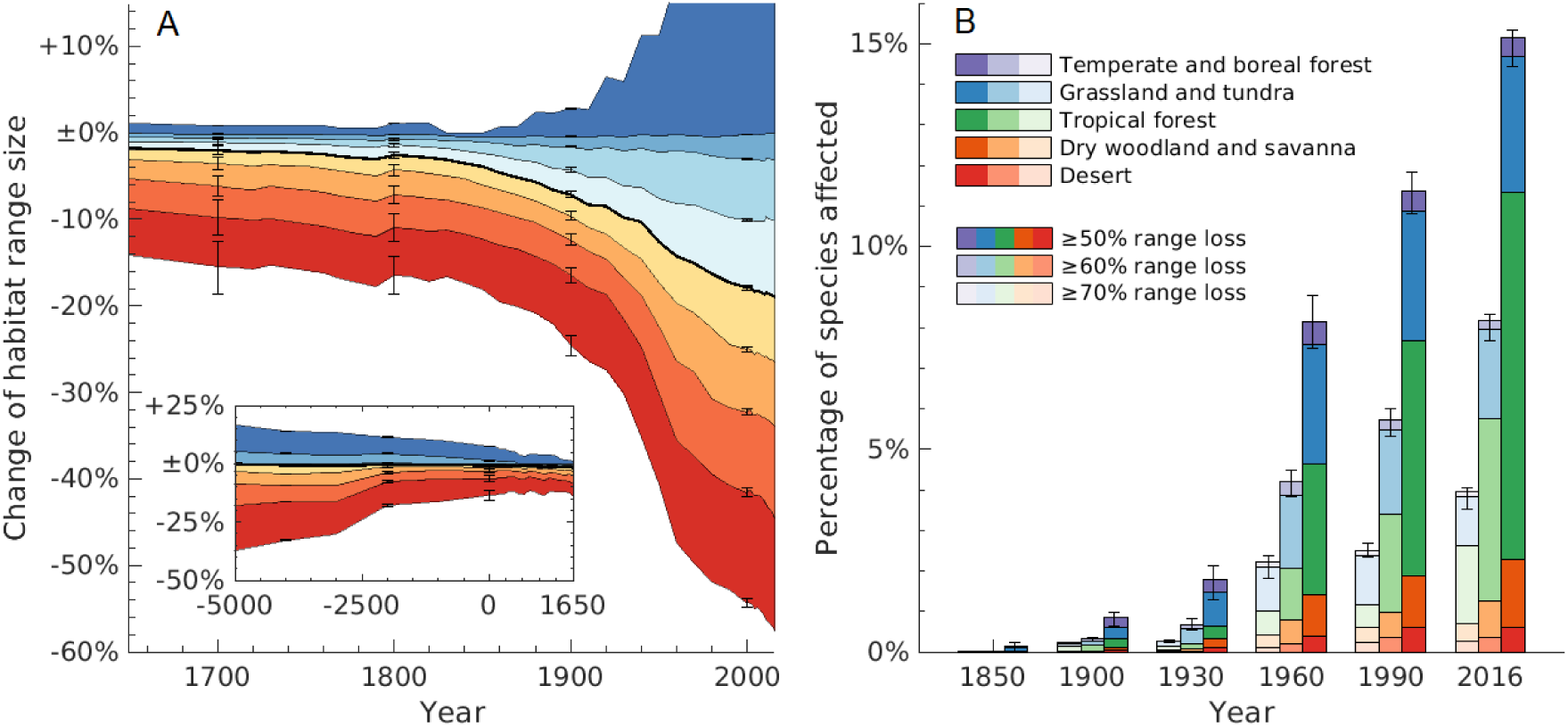
Past changes in the range sizes of mammals, birds, and amphibians. **(A)** Estimated changes in range sizes between 5,000 BCE and 2016 CE, relative to potential natural ranges in 1850 (Methods). Colour shades represent 10%, 20%, …, 90% percentiles (red, orange, …, blue) of range changes across the 16,511 species considered; the black line shows the median. **(B)** Percentages of species affected by critical range contractions, broken down by species’ primary mega-biome (Methods). Uncertainties of the quantiles in (A) and the critical losses in were derived by repeating the analysis using the available upper and lower uncertainty bounds of the land use reconstruction (Methods).

The magnitude of range contractions estimated for the past 7,000 years is not merely the result of the increasing area of converted land. Over recent centuries and millennia, species habitat loss has increased quadratically in relation to the total size of agricultural and urban areas (Fig. 2A). Whilst the first billion hectares (converted until c. 1720) caused an average 1% range loss, the most recently converted half billion hectares (since c. 1950) are responsible for an average loss of 8% of natural range sizes. This trend is mirrored by a long-term increase in the marginal impact of newly converted areas on species ranges. The average loss of habitat caused by a unit of land converted in the 20th century was 143% higher than in the 19th century, and 245% higher than in the 18th century.

**Fig. 2.**
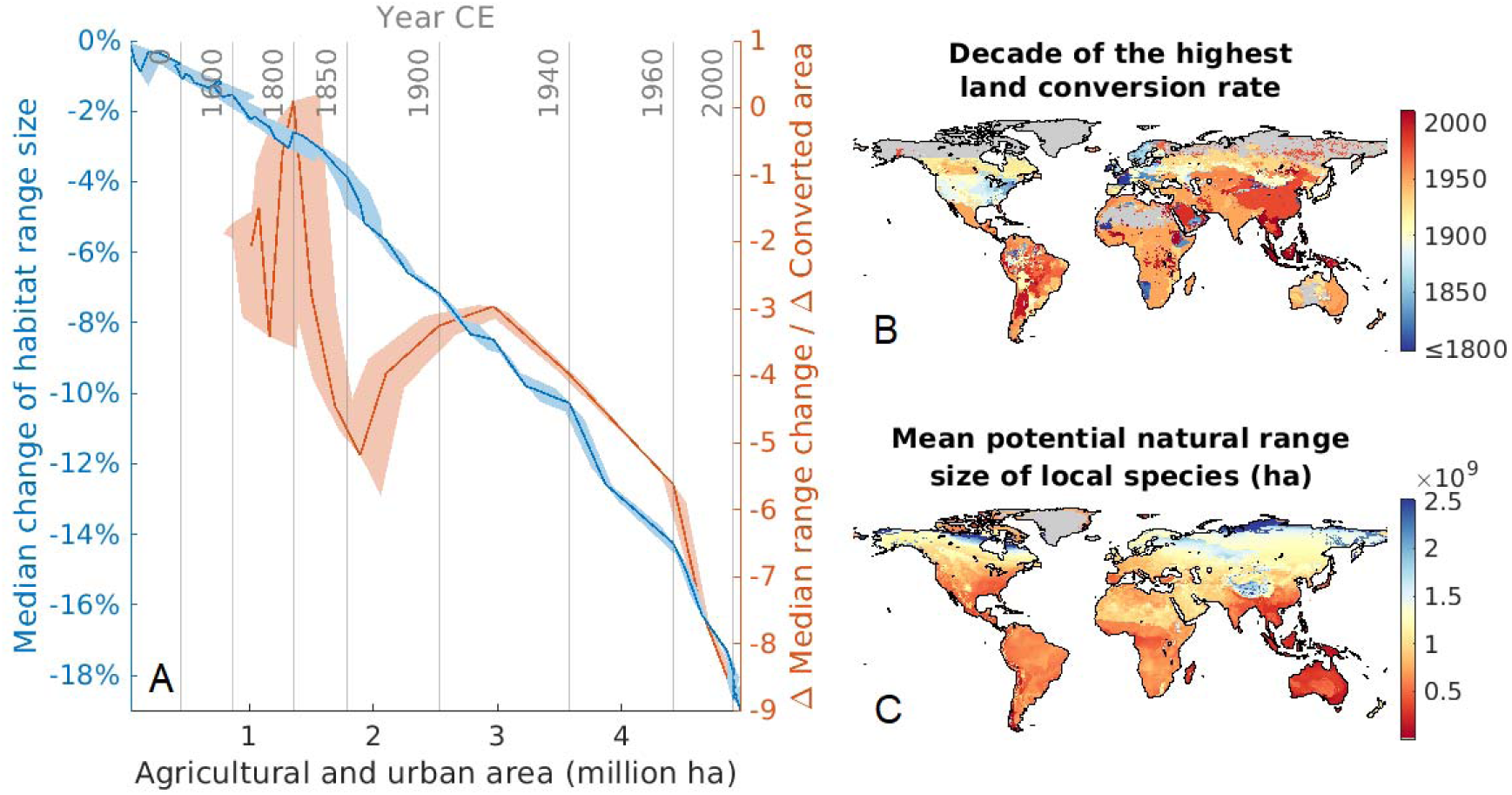
Acceleration of the marginal impact of land use on species’ range sizes. **(A)** Median range loss across all species against the cumulative global agricultural and urban area (blue line), and its derivative, the marginal impact of 1 ha of converted land on average species range loss (orange line; restricted to times for which there is at least decadal land use data). Uncertainty bands are based on the uncertainty of the land use reconstruction. **(B)** Mean potential natural range size of species locally present for the natural biome distribution in 1850 and no land use. **(C)** Local time of the highest rate of land conversion up until 2010.

This acceleration of marginal range losses can be explained by a long-term trend in the location of land use change towards tropical regions, where both local species richness is higher and average species ranges are smaller, and thus where the destruction of natural habitat leads to particularly high relative range losses ^35^ (Fig. 2B). After a long period of much less land conversion than in other parts of the world, these areas have experienced a rapid expansion of agriculture since the end of the 19th century. Habitat conversion rates reached their highest levels to date in South America around the mid-late 20th century, and in the late 20th and early 21st century in South East Asia, a global hotspot of small-ranged species ^35^ (Fig. 2C).

Whether these past trends in range losses will reverse, continue, or accelerate will depend on the global emission and socio-economic pathway chosen in the coming years and decades. By 2100, average range losses could reach up to 32% in the worst case scenario (RCP 6.0, SSP 3), or drop to 15% – roughly equivalent to levels in 1970 – in the best case (RCP 2.6, SSP 1) (Fig. 3). The proportion of species suffering the loss of at least half of their natural range could reach between 18% (RCP 2.6, SSP 1) and 35% (RCP 6.5, SSP 3) by 2100 (Fig. S3B). Isolating the impact of climate change shows that increasing levels of global warming result in more species experiencing critical range losses, while others expand their ranges ^10,13^ (Fig. S1). For any given socio-economic pathway, increasing emission levels consistently result in higher average range losses by 2100 (Fig. 3). At the same time, the differences between climate change scenarios, in terms of average range change, are sometimes smaller than the differences between socio-economic scenarios (e.g. average range loss associated with RCP 8.5/SSP 5 is smaller than that for RCP 4.5/SSP 3). Across climate change scenarios, average range loss is consistently lowest for SSP 1, similar for SSPs 2,4,5, and highest for SSP 3. While SSP 1 would enable the re-expansion of ranges in many parts of the world as the result of the abandonment of agricultural areas, notably in Southeast Asia, SSP 3 represents a continuation of land use change in the tropics, most strongly in the Congo basin (Fig. S4).

**Fig. 3.**
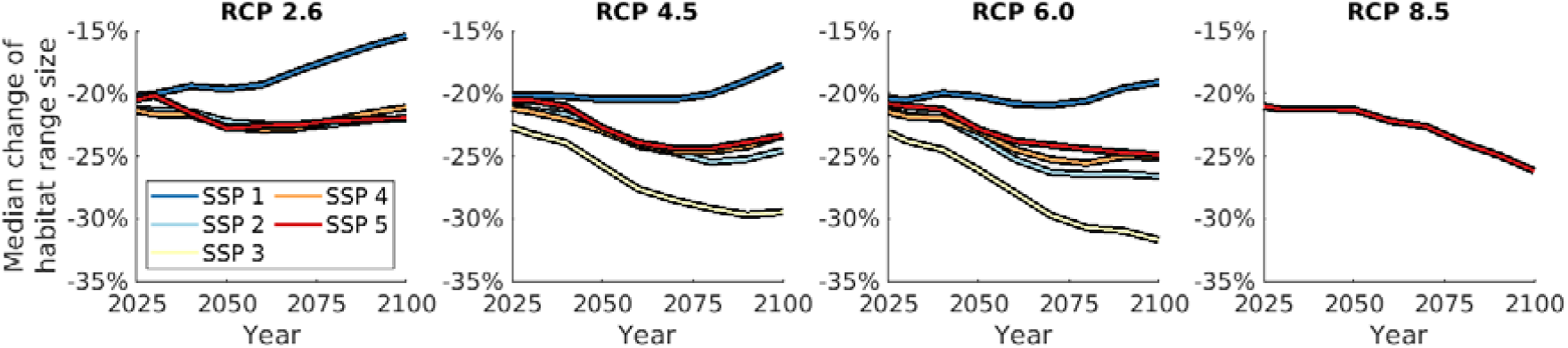
Future projections of median range change across species (analogous to the black line in Fig. 1A) until the year 2100 for RCPs 2.6, 4.5, 6.0 and 8.5, and SSPs 1–5. Complete fan charts showing 10–90% percentiles of range changes across species (as in Fig. 1A) for individual RCP/SSP combinations are available as Fig. S3A.

Our estimates of the past and current states of species habitat ranges, and how they will be impacted under alternative future climatic and socio-economic scenarios, provide important evidence for conservation-oriented decision-making from the local to the global scale. Our results provide quantitative support for policy measures aiming at curtailing the global area of agricultural land (by closing yield gaps, encouraging dietary shifts, and stabilising population growth) ^36^, especially in areas of small-range species ^35^, steering production to agro-ecologically optimal areas when additional expansion is inevitable ^37,38^, targeting land abandonment and restoration in hotspot areas ^39,40^, and limiting climate change ^41^. Whilst our data quantify the drastic consequences for species ranges if global land use and climate change are left unchecked, they also demonstrate the tremendous potential of timely and concerted policy action for halting and indeed partially reversing previous trends in global range contractions.

## Acknowledgments

The authors are grateful to BirdLife International for sharing and advising on the bird species distribution data, and to Mario Krapp for advice on the climatic data.

## Author contributions

R.M.B. conducted the analysis and wrote the manuscript. All authors designed the project, interpreted the results, and revised the manuscript.

## Methods

Our reconstructions and projections of species range sizes use maps of the global distribution of cropland, pasture, and built-up land, and natural biomes, through time. Here, we first describe how these input data were derived, and then specify how they were combined with species distribution and habitat data to estimate range sizes.

### Global land use data

For the time period 5,000 BCE – 2016 CE, we used reconstructions of global cropland, pasture and urban areas from the Hyde v3.2 dataset ^42^ (available from: https://doi.org/10.17026/dans-25g-gez3). Although Hyde 3.2 provides land use data as far back as 10,000 BCE, we restricted our analysis to the past 7,000 years, during which the global distribution of land area has changed only very little, at the spatial resolution of our analysis. Prior to 7,000 years ago, significantly lower global sea levels ^43^ resulted in the emergence of new land areas, which our estimation of species’ ranges does not accommodate. Archaeological data has recently been argued to suggest underestimations of land use in some regions before 1850 CE in the Hyde dataset ^44^. With this potential caveat in mind, the Hyde dataset remains the most widely used spatially-explicit long-term reconstruction of global land use ^45^. A total of 69 maps, including lower and upper uncertainty bounds, are available at 1000-year time steps between 5,000 BCE and 0 CE, then at 100-year time steps until 1700 CE, at 10-year until 2000 CE, and at 1-year time steps until 2016 CE. These data were upscaled from their original spatial resolution of 0.083° to a 0.5° grid.

For the period 2020–2100 CE, we used 0.5°-resolution 10-year-time-step projections of global cropland, pasture and urban areas from the AIM model ^45^ (available from: https://doi.org/10.7910/DVN/4NVGWA), covering Representative Concentration Pathways (RCPs) 2.6, 4.5, 6.0 and 8.5, and Shared Socio-economic Pathways (SSPs) 1–5. The dataset contains all possible combinations of these emission and socio-economic trajectories with the exception of RCP 2.6/SSP 3, and RCP 8.5/SSPs 1–4. We refer to refs. ^27–29,46^ for details of the emission and socio-economic pathways, and to ref. ^28^ for a comparison between the AIM model and other Integrated Assessment Models.

### Global biome data

We used the Biome4 vegetation model ^47,48^ (available from: https://pmip2.lsce.ipsl.fr/synth/biome4.shtml) to simulate the distribution of global potential natural biomes between 5,000 BCE and 2016 CE, and between 2020 and 2100 CE for each of the four climate change scenarios considered (RCPs 2.6, 4.5, 6.0, 8.5), at a spatial resolution of 0.5°. Inputs required by Biome4 include global mean atmospheric CO_2_ concentration, and gridded monthly means of temperature, precipitation, and percent sunshine. Past and RCP-specific future CO_2_ levels were obtained from refs. ^49–51^ and ref. ^52^, respectively. The climatic data were generated as follows: For the period 5,000 BCE – 1900 CE, we used a dataset of 0.5° downscaled and bias-corrected climate reconstructions ^53^ (available from: https://doi.org/10.6084/m9.figshare.12293345.v3), which is based on HadCM3 and HadAM3H climate simulations ^54^. For the period 1900–2016 CE, we used 0.5°-resolution annual global observational data ^55^ (available from: https://doi.org/10.5285/10d3e3640f004c578403419aac167d82). For the period 2020–2100 CE, and for each RCP (2.6, 4.5, 6.0, 8.5), we used annual 1.875°×1.25°-resolution HadGEM2-ES simulations ^56^, averaged across available ensembles r1i1p1, r2i1p1, r3i1p1, r4i1p1 (available from: https://esgf-node.llnl.gov/search/cmip5/), which we downscaled and bias-corrected by means of the Delta Method ^57^ and 2006 observed climate ^58^. By construction of the Delta Method, the final time slice of the downscaled and bias-corrected past simulations, and the first time slice of future simulations, are identical to the observational climate at the appropriate time; thus, our full set of past and future climatic data are continuous in time.

For the computation of the global biome distribution at a point in time *T*, we used as inputs for Biome4 the CO_2_ value and gridded monthly climate values averaged across the time interval [*T* – 30 years, *T*]. Pre-1900 climatic data was temporally interpolated to the relevant years where necessary. We performed biome simulations in 1,000-year time steps between 5,000 BCE and 0 CE, then in 200-year time steps until 1400 CE, in 100-year time steps until 1700 CE, in ∼20-year time steps until 1930, and in ∼15-year time steps until 2016 CE, as well as between 2020 and 2100 CE. Biome4 estimates ice biomes only based on climatic conditions, which can lead to the spatial extent of ice sheets being underestimated. We therefore changed simulated non-ice biomes to ice in grid cells covered by ice sheets according to the ICE-7g dataset ^59^ at the relevant times. The complete time series of global biome simulations is available as Supplementary Movie 1.

### Estimation of species ranges

We estimated the geographic ranges of individual bird, mammal and amphibian species through time following the general methodology in ref. ^22^, while considering a much larger set of species, points in time, and land use and climatic scenarios. The approach combines the following data:

I. Spatial polygon data on species-specific extents of occurrence of all known birds ^60^ (available from: http://datazone.birdlife.org/species/requestdis), mammals and amphibians ^61^ (available from: https://www.iucnredlist.org/)
II. Species-specific biome types required for suitable habitat ^43,44^ (data available from the same websites)
III. Maps of global potential natural biome distributions corresponding to the relevant climatic conditions over time (i.e. reconstructions for the past, and RCP-specific projections for the future)
IV. Maps of global cropland, pasture and built-up land through time (i.e. reconstructions for the past, and RCP- and SSP -specific projections for the future)

The data I–IV were used to estimate the range of individual species at a given point in time as follows. First, we used species-specific extents of occurrence (data I), which we remapped from their original spatial polygon format to a 0.5° resolution grid. These data provide rough spatial envelopes of each species’ maximum geographic range extent, which do not account for the distribution of natural or artificial land cover within that area, and therefore extend substantially beyond a species’ actual area of occupancy ^62,63^. We refined these data by combining species-specific biome requirements (data II) with maps of the global biome distribution at a given point in time (data III; see section above). This required matching IUCN habitat categories (https://www.iucnredlist.org/resources/habitat-classification-scheme) with the biome categories of the Biome4 vegetation model, which was done as shown in Table S1. Thus, we subset the previous spatial data by only retaining grid cells where the natural biome type is included in the species’ list of suitable habitat categories. The result of this step represents a species’ estimated potential natural habitat range (i.e. in the hypothetical absence of anthropogenic land use) at a given point in time. In order to estimate actual habitat ranges, we used maps of global land use through time (data IV; see section above), and removed the proportion of cropland, pasture and urban land in each grid cell contained in the potential natural range at the relevant point in time, unless the species’ list of suitable habitat categories (data II) contained the relevant land cover type. Numerical range sizes were computed as the sums of the remaining areas of the grid cells that form the ranges.

We applied this method at each point in time for which global land use data is available (see section above). In this way, we obtained potential natural ranges and actual ranges for 69 points in time between 5,000 BCE and 2016 CE, and for 8 points in time between 2020 and 2100 CE, for each of which ranges were derived for 16 combinations of future climatic and socio-economic pathways (see section above). We repeated this analysis using the lower and upper uncertainty bounds of the Hyde 3.2 land use reconstructions, which allowed us to quantify the uncertainties of our estimates of past range size changes with respect to the underlying land use data.

Since the global distribution of biomes changes over time as the result of (naturally or anthropogenically) changing climatic conditions, the estimated potential natural range sizes are time-dependent. This motivates to consider range changes in relation to the potential natural ranges estimated at a reference time, for which we chose the year *t*_*0*_ = 1850, representing a modern pre-industrial baseline.

Denoting the potential natural range and the actual range of a species *i* at a time *t* by 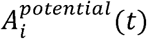 and 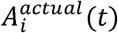, respectively, the range change associated with species *i* at time *t* as the result of the distribution of biomes and land use at that time was computed at as

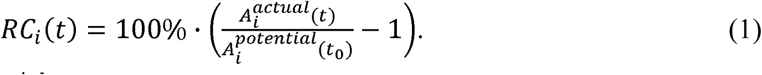

Species for which 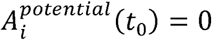, i.e. for which changes in range size cannot be computed, were removed from the analysis. This can occur even when a species’ true potential natural range size at *t*_*0*_ is not zero if there is a mismatch within the species’ extent of occurrence between the true biome distribution and the one simulated by Biome4. Other than due to flaws in the climate data or Biome4, this can be a consequence of the spatial resolution of the vegetation data, when the relevant 0.5° grid cells contain microclimate-specific biomes that are different from ones corresponding to the cell-average climates, simulated here.

Based on the set {*RC*_*i*_ (t)}_*i*=1,2,…_ of the individual range changes of all species through time, we calculated range change percentiles at each point in time (Fig. 1A), and determined the proportion of species that have experienced the loss of a given percentage of their baseline range (Fig. 1B). Similarly as in Eq. (1), we also computed the range change attributed only to climate-change-induced biome changes, 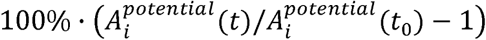 (Fig. S1). For each species *i*, point in time *t* and, in the case of future projections, RCP and SSP scenario, potential and actual ranges 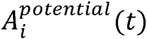 and 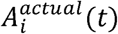 are available as Supplementary File S1.

Our method of estimating species’ spatial ranges through time shares the advantages and limitations of the original approach presented in ref. ^22^. As the empirical data on species habitat preferences only provide categorical biome requirements, not continuous climatic envelopes, the method does not account for range changes resulting from changes in climatic conditions that are too small to manifest as biome changes. However, estimating exact climatic envelopes of species can be subject to substantial uncertainty, and be highly sensitive to the way in which they are estimated (see below). By construction of the method, species’ ranges over time vary within the extents of occurrence provided with the empirical data, and do not exceed those. Justification for this assumption is provided by fact that potential natural ranges (and much more, actual ranges) are generally well-contained within extents of occurrence, with the former accounting for an average of 59% of the area of the latter in 1850, thus providing ample space for range expansions and shifts within the boundaries. Additional evidence that the restriction of ranges to the extents of occurrence does not prevent significant range expansions can be seen in the sizeable number of species that are projected to experience such range expansions in future scenarios of strong global warming (Fig. S1).

Climate niche models estimate statistical relationships between climatic conditions and species’ spatial distributions, and apply these to climate projections to estimate future distribution patterns ^64^; as such, they are not subject to the above limitations. However, they suffer from other important drawbacks that challenge their suitability for our specific approach. Previous evaluations of niche-based simulations have pointed out issues related to a lack of adequate data for model calibration ^65,66^, risks of parameter overfitting ^67^, unreliable behaviour when extrapolating beyond current conditions ^68^, high sensitivity to the assumed statistical model ^69^, mismatches with empirically observed ranges ^70^, lack of explanatory power ^71^, and methodological issues ^72^, all of which can result in considerable uncertainties in the simulated species distributions ^64–75^. These issues would make it difficult to ensure the level of robustness that is necessary in an analysis involving the large number of species, times, and scenarios considered here. By operating directly on the empirical data on species’ extents of occurrence and biome requirements, and not being reliant on any particular statistical model or parameterisation, the approach used here is particularly transparent ^22,76^ and provides the necessary robustness.

## Supplementary Figures

**Fig. S1.**
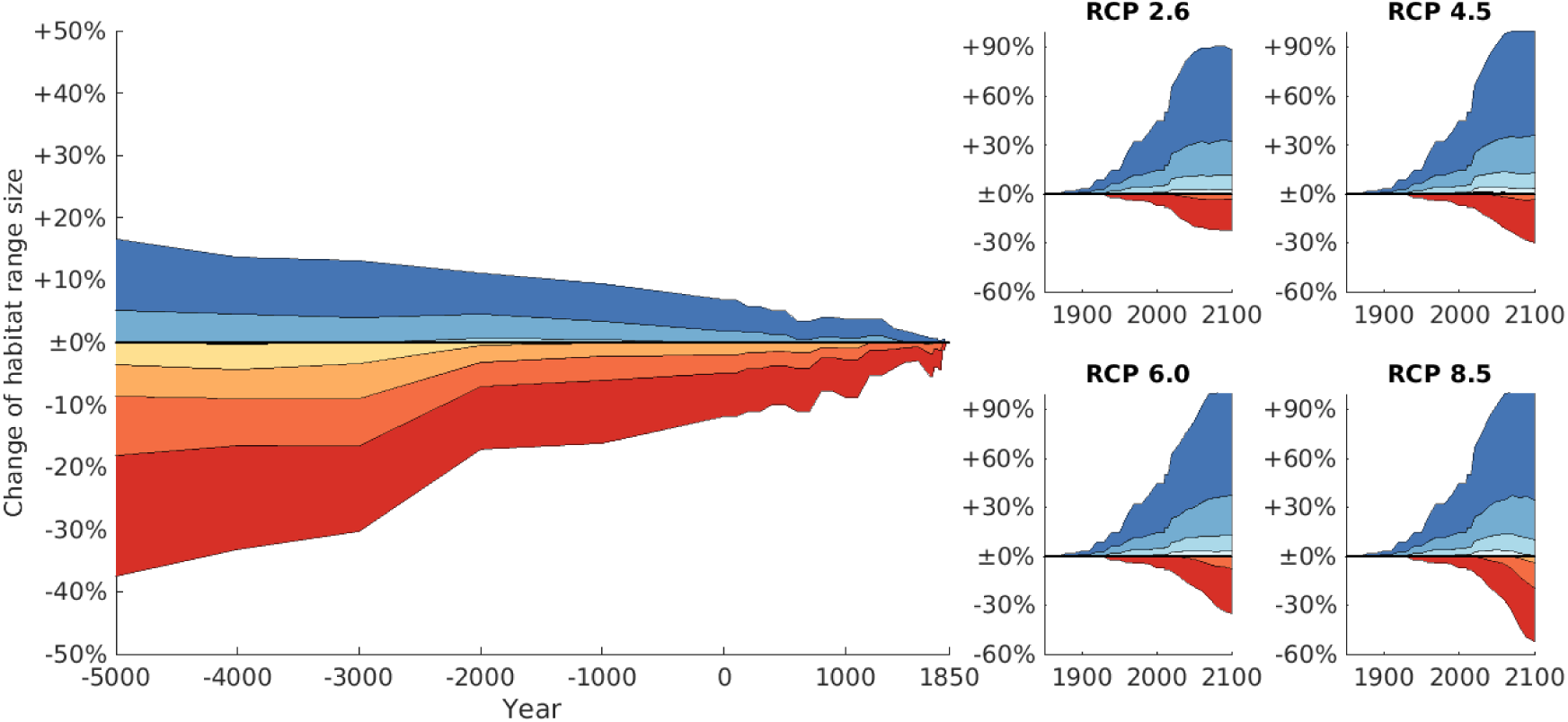
Past and RCP-specific future changes in range sizes through time based only on climate-driven biome distributions (i.e. in the hypothetical absence of anthropogenic land use), relative to potential natural range sizes in 1850. Colour shades and the black line represent 10–90% percentiles and the median of range changes across all species, respectively (analogous to Fig. 1A).

**Fig. S2.**
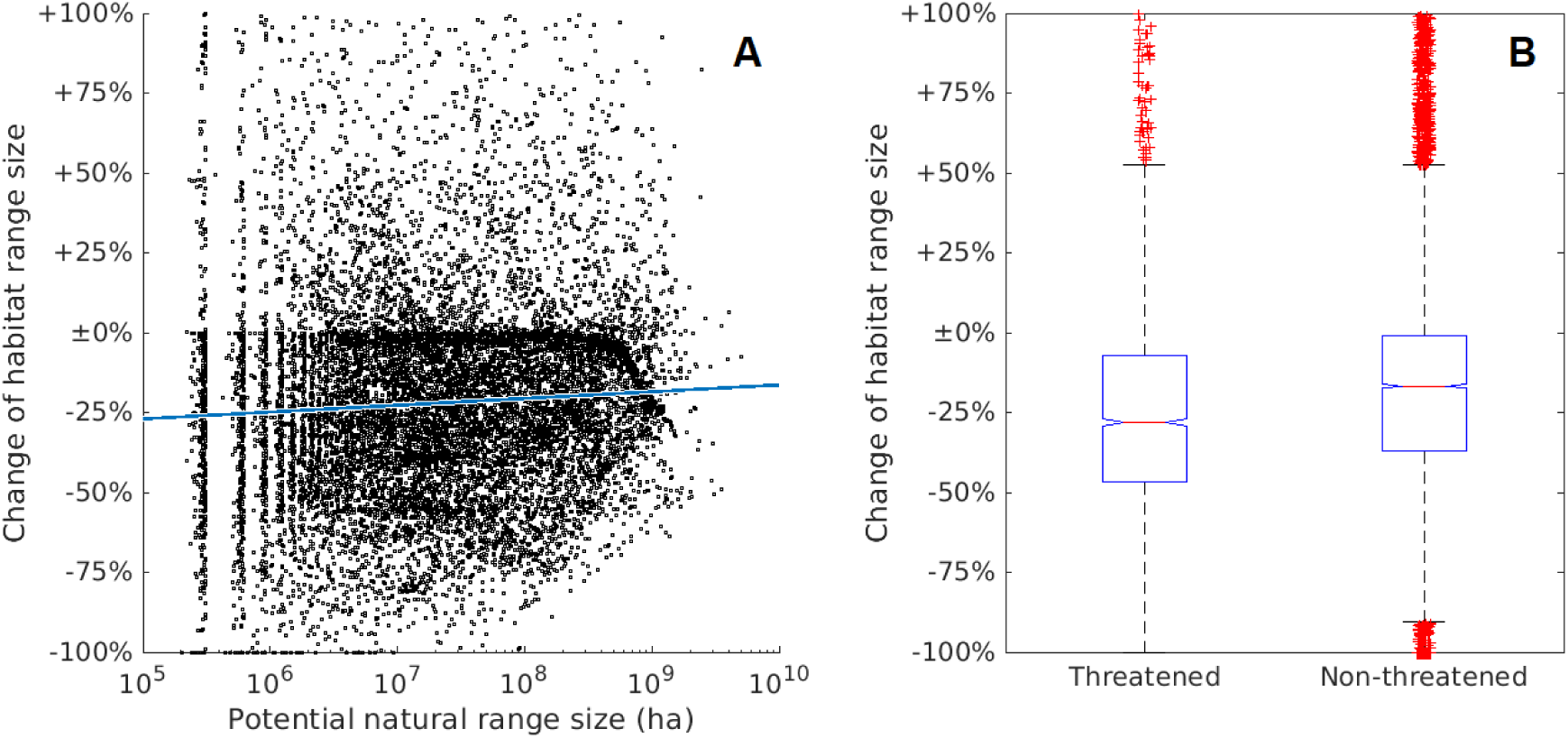
Range changes in small-ranged and threatened species. **(A)** Changes of range sizes in 2016 (relative to potential natural range sizes in 1850) against potential natural range size. The log-linear relationship (red line) does not explain much variation of the data (R^2^ < 0.1) but is significant (p < 0.01). **(B)** Change of range sizes in 2016 of threatened and non-threatened species. Threatened species include those classified as vulnerable, endangered and critically endangered ^60,61^. Red lines represent medians, blue box limits represent upper and lower quartiles, whiskers represent 1.5 × interquartile range, red crosses represent outliers. Median range loss for threatened species is significantly higher than for non-threatened species (p < 0.01). For visualisation purposes, y-axes were capped at +100% range size increases.

**Fig. S3.**
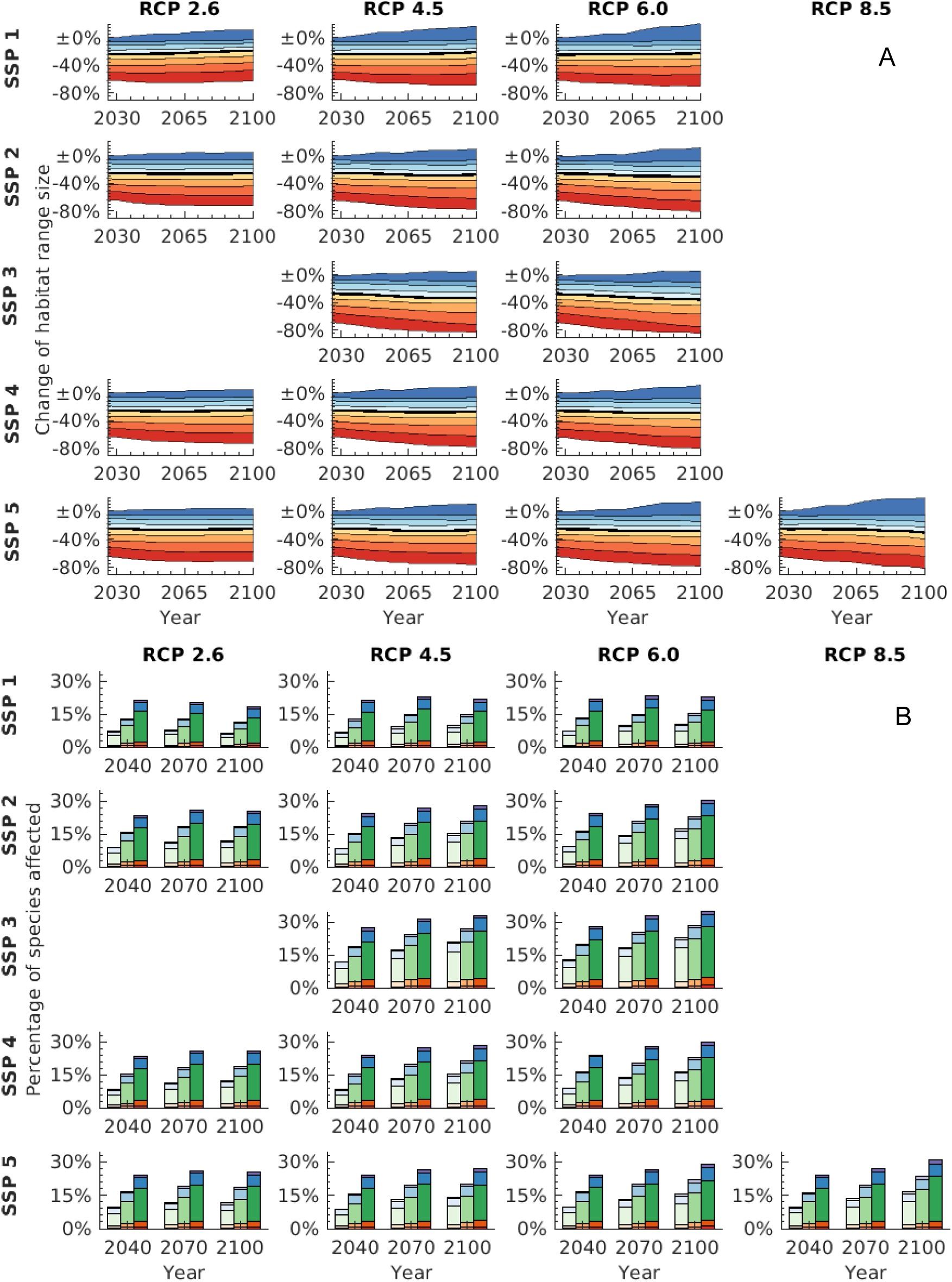
Projected future changes in species’ range sizes. **(A)** Future SSP- and RCP-specific changes in the habitat range sizes between 2025 and 2100 CE, relative to potential natural range sizes in 1850. Colour shades and the black line represent 10–90% percentiles and the median of range changes across all species, respectively. **(B)** Percentages of species affected by critical range losses, broken down by species’ primary mega-biome. Colour schemes are identical to those in Fig. 1.

**Fig. S4.**
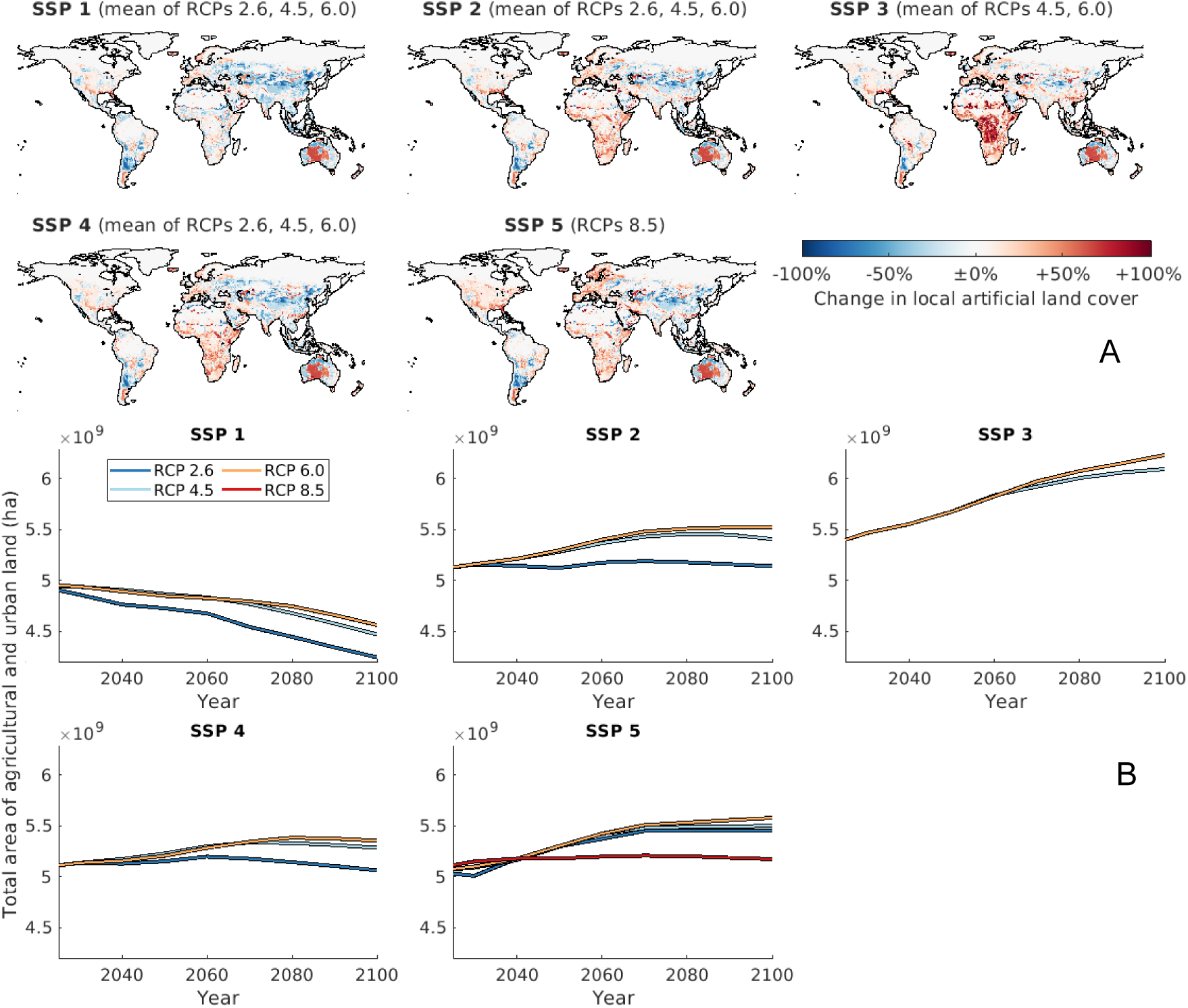
Overview of future land use scenarios. **(A)** Difference between SSP-specific distributions of agricultural and urban land in 2100 (averaged across the RCPs available for the relevant SSP) and the current (year 2016) distribution. **(B)** SSP- and RCP-specific trajectories of the total area of converted land between 2025 and 2100.

## Supplementary Tables

**Table S1.**
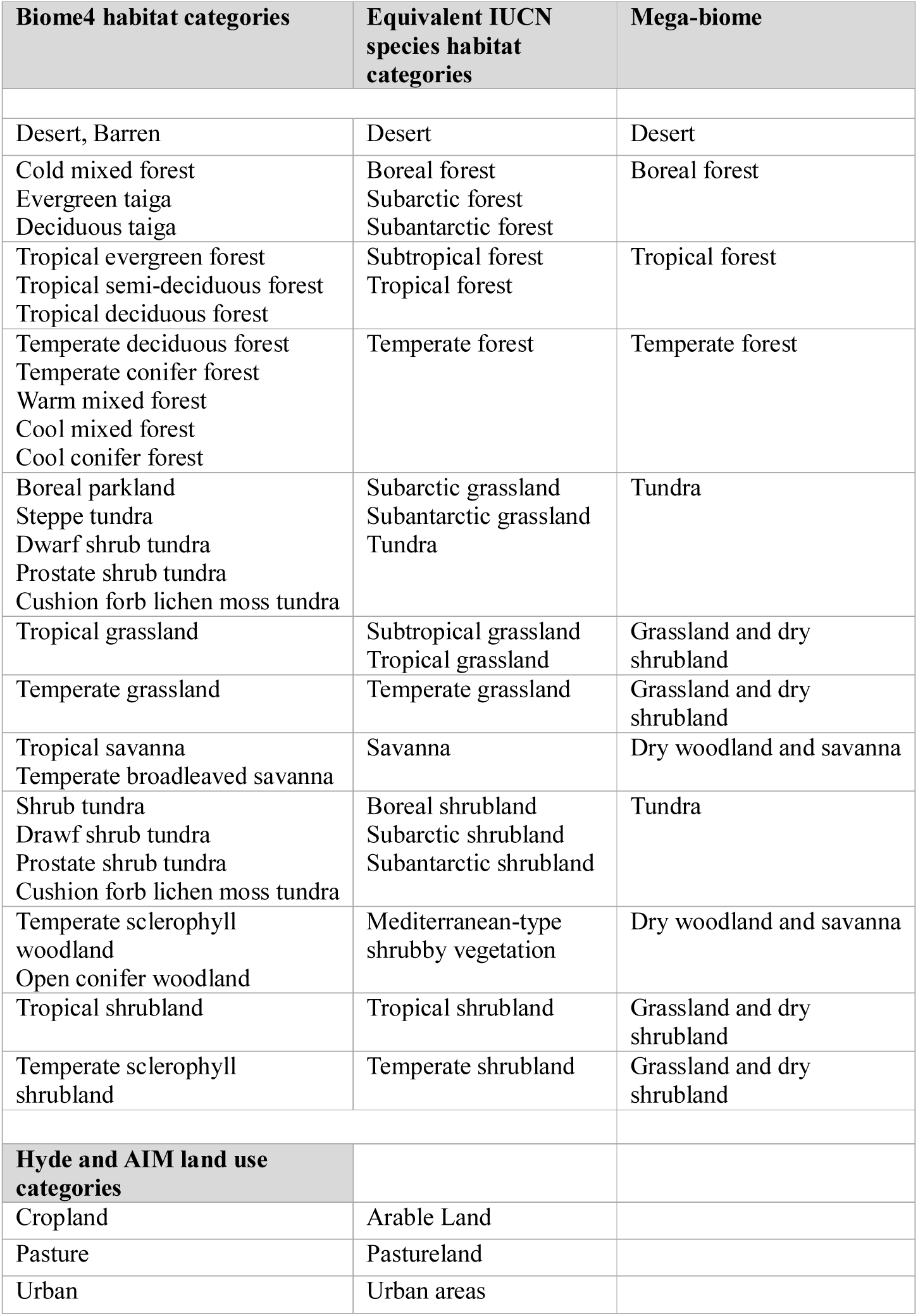
Matching between land cover types in the global land use and Biome4 data, and IUCN categories of species habitats. Species-specific primary mega-biomes were assigned based on the mega-biome shown in the third column that accounts for the largest area of the species’ potential natural range in 1850.

## Supplementary Movies

### Movie S1

Time series of past and RCP-specific future global distributions of biomes.

## Supplementary Files

### File S1

Past and RCP/SSP-specific future potential natural and actual ranges of 16,511 species through time.

